## supplemental figures S1 S2 S3 for "Adaptation of a commercial NAD quantification kit to assay the base exchange activity of SARM1"

**SUPPLEMENTAL INFORMATION**

(associated to Figure 2 & Figure 3)

**Fig S1**

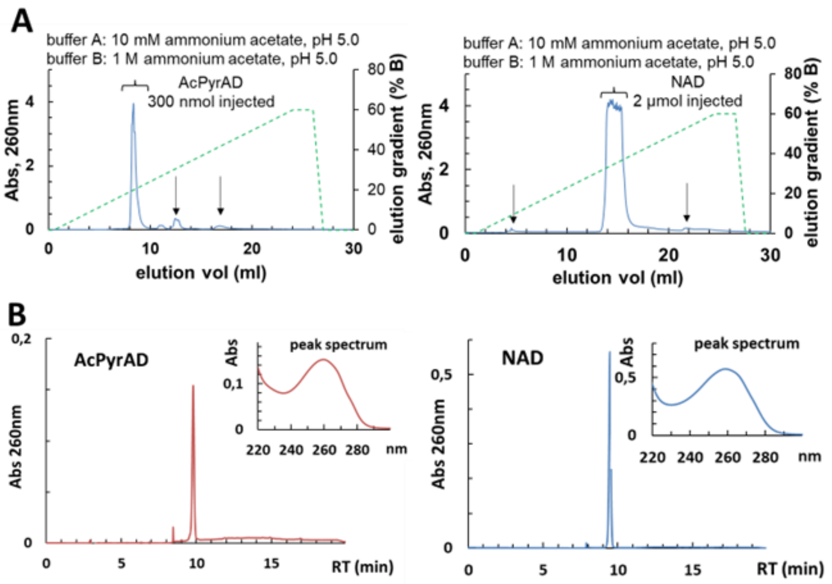

**Figure S1. IEC-FPLC purification of commercial stocks of either NAD or AcPyrAD**

(**A**) Typical chromatographies of AcPyrAD (left) or NAD (right) carried out for cleaning. Purchased dinucleotides were dissolved in water and injected on TSK-DEAE (Tosoh column, 250x4.6mm) equilibrated in buffer A, followed by elution at 1 ml/min by a gradient up to 60% of buffer B as indicated. Peaks below parentheses were collected and lyophilized. Black arrows indicate the most frequent contaminants removed.

(**B**) C18-HPLC analysis of the two purified compounds resuspended in milliQ water. Insets, UV scan profiles of the single peaks eluted.

(associated to Figure 4)

**Fig S2**

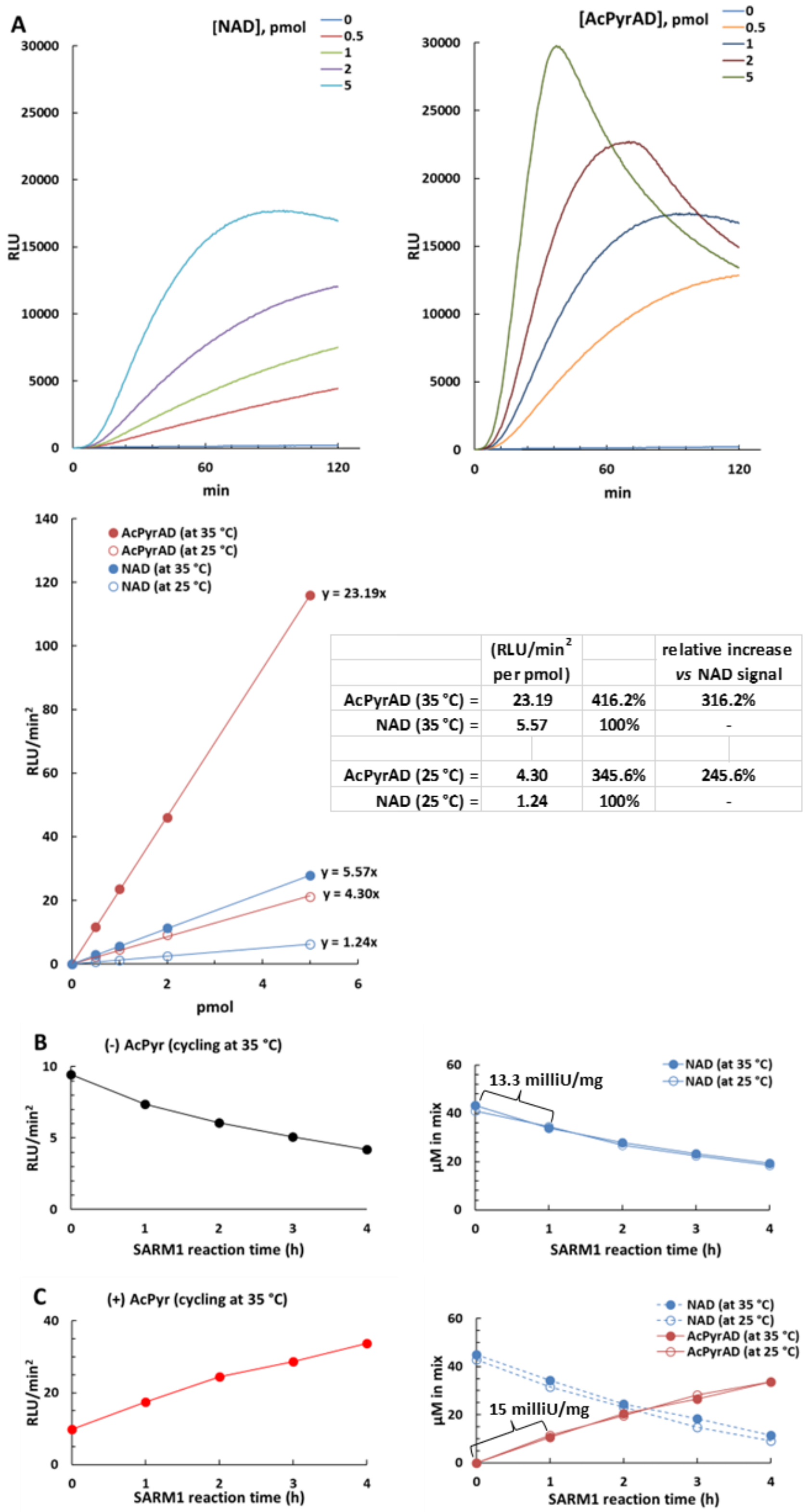

| | (RLU/min² per pmol) | | relative increase *vs* NAD signal |
|---|---|---|---|
| AcPyrAD (35 °C) = | 23.19 | 416.2% | 316.2% |
| NAD (35 °C) = | 5.57 | 100% | - |
| AcPyrAD (25 °C) = | 4.30 | 345.6% | 245.6% |
| NAD (25 °C) = | 1.24 | 100% | - |

**Figure S2. Cycling amplification of NAD or AcPyrAD at 35 °C & re-building at this higher temperature of data in Fig 4**

(**A**) Luminescence emission (RLU) from 0 to 5 pmol of each NAD and AcPyrAD after cycling at 35 °C, followed by RLU/min$^2$ calculation and plotting as above. The cycling slopes obtained at 35 °C for these two standards resulted higher than previous values at 25 °C (see table data and both Fig 2A & Fig 3A of main text). Nonetheless, the relative increase of luminescence from AcPyrAD referred to that from NAD at each temperature is largely equivalent. Linearization showed $R^2$ values ≥ 0.99 in all cases.

(**B** and **C**) SARM1 reactions (already analyzed, see Fig 4, main text) were re-evaluated through the NAD/NADH-Glo™ Assay at 35 °C. Data are color-coded like in Figure 4A. The results are superimposable to those obtained previously at 25 °C (open circles).

(associated to Figure 4)

**Fig S3**

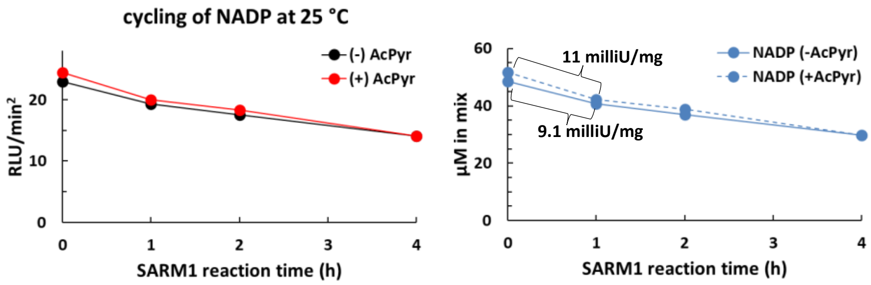

| CYCLING SLOPE of STANDARD | |
|---|---|
| (RLU/min$^2$ per pmol) | |
| NADP (A) = | 11.841 |

**Figure S3 – A typical reaction of AcPyr base exchange on NADP catalysed by SARM1 evaluated by NADP/NADPH-Glo™ Assay at 25 °C**

Reaction mixtures contained recombinant SARM1 and 50 µM NADP as substrate, plus or minus 2 mM AcPyr. They were treated and processed as the mixtures with NAD (see Fig 4, main text) but cycled through the NADP/NADPH-Glo™ Assay at 25 °C together with a NADP standard. Luminescence emitted (RLU/min$^2$ values) was used to calculate the levels of NADP in the control minus AcPyr (continuous blue line) or in the sample plus AcPyr (dotted blue line). Calculations were made using equation 1 in Methods and the standard slope A for NADP as indicated. NADP consumption rates were calculated within the first one hour of incubation.
